# Stochastic models of gene transcription with upstream drives: exact solution and sample path characterization

**DOI:** 10.1101/055202

**Authors:** Justine Dattani, Mauricio Barahona

## Abstract

Gene transcription is a highly stochastic and dynamic process. As a result, the mRNA copy number of a given gene is heterogeneous both between cells and across time. We present a framework to model gene transcription in populations of cells with time-varying (stochastic or deterministic) transcription and degradation rates. Such rates can be understood as upstream cellular drives representing the effect of different aspects of the cellular environment. We show that the full solution of the master equation contains two components: a model-specific, upstream effective drive, which encapsulates the effect of the cellular drives (e.g., entrainment, periodicity or promoter randomness), and a downstream transcriptional Poissonian part, which is common to all models. Our analytical framework allows us to treat cell-to-cell and dynamic variability consistently, unifying several approaches in the literature. We apply the obtained solution to characterize several gene transcription models of experimental relevance, and to explain the influence on gene transcription of synchrony, stationarity, ergodicity, as well as the effect of time-scales and other dynamic characteristics of drives. We also show how the solution can be applied to the analysis of single-cell data, and to reduce the computational cost of sampling solutions via stochastic simulation.

## I. INTRODUCTION

Gene transcription, the cellular mechanism through which DNA is copied into mRNA transcripts, is a complex, stochastic process [1]. As a result, the number of mRNA copies for most genes is highly heterogeneous both within each cell over time, and across cells in a population [2–4]. Such fundamental randomness is biologically relevant: it underpins the cell-to-cell variability linked with phenotypic outcomes and cell decisions [5–9].

The full mathematical analysis of gene expression variability requires the solution of *master equations*. Given a gene transcription model, its master equation (ME) is a differential-difference equation that describes the evolution of *P*(*n,t*), the probability of having *n* mRNA molecules in a single cell at time *t*. However, MEs are problematic to solve, both analytically and numerically, due to the difficulties associated with discrete stochastic variables—the molecule number *n* is an integer [10]. Indeed, most existing analytical solutions of the ME are specific to particular models, typically obtained via the probability generating function and under stationarity assumptions [11–17]. When analytical solutions are intractable, the first few moments of the distribution are approximated, usually at stationarity, but error bounds are difficult to obtain [18, 19]. Alternatively, full stochastic simulations are used, although the computational cost to sample *P*(*n*, *t*) at each t is often impractical, and many methods lead to estimation bias in practice [20].

With the emergence of accurate measurements of time-courses of mRNA counts in single cells [3, 4, 21–23], the application of MEs to data analysis faces new challenges. Mathematically, ME models must be able to describe time-dependent gene transcription in single cells within a population, allowing for a degree of synchrony or cell-to-cell correlation. However, current stationary solutions tacitly assume that gene expression is uncorrelated between cells; hence most current analytical models cannot account for the dynamic variability due to upstream biological drives, such as chromatin remodeling [24], transcription factor binding [25], circadian rhythms and cell cycle [26, 27], external signaling [28], or stimulus-induced modulation or entrainment [29, 30]. Full solutions of the ME that capture temporal heterogeneity from the singlecell to the population level could help unravel how the dynamics of biological phenomena affect gene transcription and regulation [31]. These examples highlight the methodological need for solvable models flexible enough for the analysis of a range of datasets, thus enabling the formulation of hypotheses in conjunction with experiments.

Here, we consider a simple, yet generic, framework for the solution of the ME of gene transcription and degradation for single cells under upstream drives, i.e., when the transcription and degradation parameters can be time-dependent functions or stochastic variables. We show that the exact solution *P*(*n*, *t*) for such a model naturally decouples into two parts: a discrete transcriptional Poisson component, which is common to all transcription models of this kind, and a model-specific continuous component, which describes the dynamics of the parameters reflecting the upstream variation. To obtain the full *P*(*n*, *t*) one only needs to calculate the probability density for the model-specific upstream drive, which we show to correspond to a continuous variable satisfying a linear random differential equation directly related to traditional differential rate equations of chemical kinetics. Below we present the properties of the general solution, including its moments and noise characteristics in terms of the moments of the upstream component. We also clarify the different effects of stochastic and deterministic drives by considering the Fano factor across the population and across time. To illustrate the utility of our approach, we present analytical and numerical analyses of several models in the literature, which are shown to correspond to different upstream drives. Finally, we provide examples of its use for data analysis.

## II. THE MASTER EQUATION FOR GENE TRANSCRIPTION IN POPULATIONS OF CELLS WITH UPSTREAM DRIVES

### Notation and formulation of the problem

To study gene expression in a single cell with time-dependent upstream drives, we consider the stochastic process in continuous time *t*, 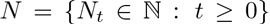, where *N_t_* is a discrete random variable describing the number of mRNA molecules in the cell. We look to obtain the probability mass function, *P*(*n*, *t*):= Pr(*N_t_* = *n*).

The mRNA copy number increases via transcription events and decreases via degradation events but, importantly, the transcription and degradation rates can depend on time and can be different for each cell (Fig. 1). To account for such variability, we describe transcription and degradation rates as stochastic processes 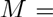 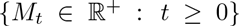 and 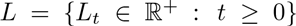, without specifying any functional form except requiring that *M* and *L* do not depend on the number of mRNA molecules already present. Deterministic time-varying transcription/degradation rates, with or without cell-to-cell correlations, are a particular case of this definition.

Following standard notation in the stochastic processes literature, *M_t_* and *L_t_* denote the random variables at time *t*. To simplify notation, however, we depart from standard notation and denote the sample paths (i.e., realisations) of *M* and *L* by {*μ*(*t*)}_*t*≥0_ and {λ(*t*)}_*t*≥0_, respectively, thinking of them as particular functions of time describing the transcription and degradation rates under the changing cellular state and environmental conditions in an ‘example’ cell (Fig. 1). The sample paths of other random variables are denoted similarly, e.g., the sample paths of *N_t_* are {*v*(*t*)}_*t*≥0_.

The sample paths {*μ*(*t*)}_*t*≥0_ and {λ(*t*)}_*t*≥0_ represent *cellular drives* encapsulating the variability across time and across the population consistently. This formulation unifies several models in the literature, which implicitly or explicitly assume time-varying transcription and/or degradation processes [4, 13, 32–36], and can be shown to correspond to particular types of dynamic upstream variability. In addition, the framework allows us to specify cell-to-cell correlations across the population, which we refer to as the ‘degree of synchrony’. A population will be *perfectly synchronous* when the sample paths of the drives for every cell in the population are identical, i.e., if *M_t_* and *L_t_* have zero variance. If, however, transcription and/or degradation rates differ between cells, *M_t_* and *L_t_* themselves emerge from a probability density: the wider the density, the more asynchronous the cellular drives are (Fig. 1).

**Fig. 1.**
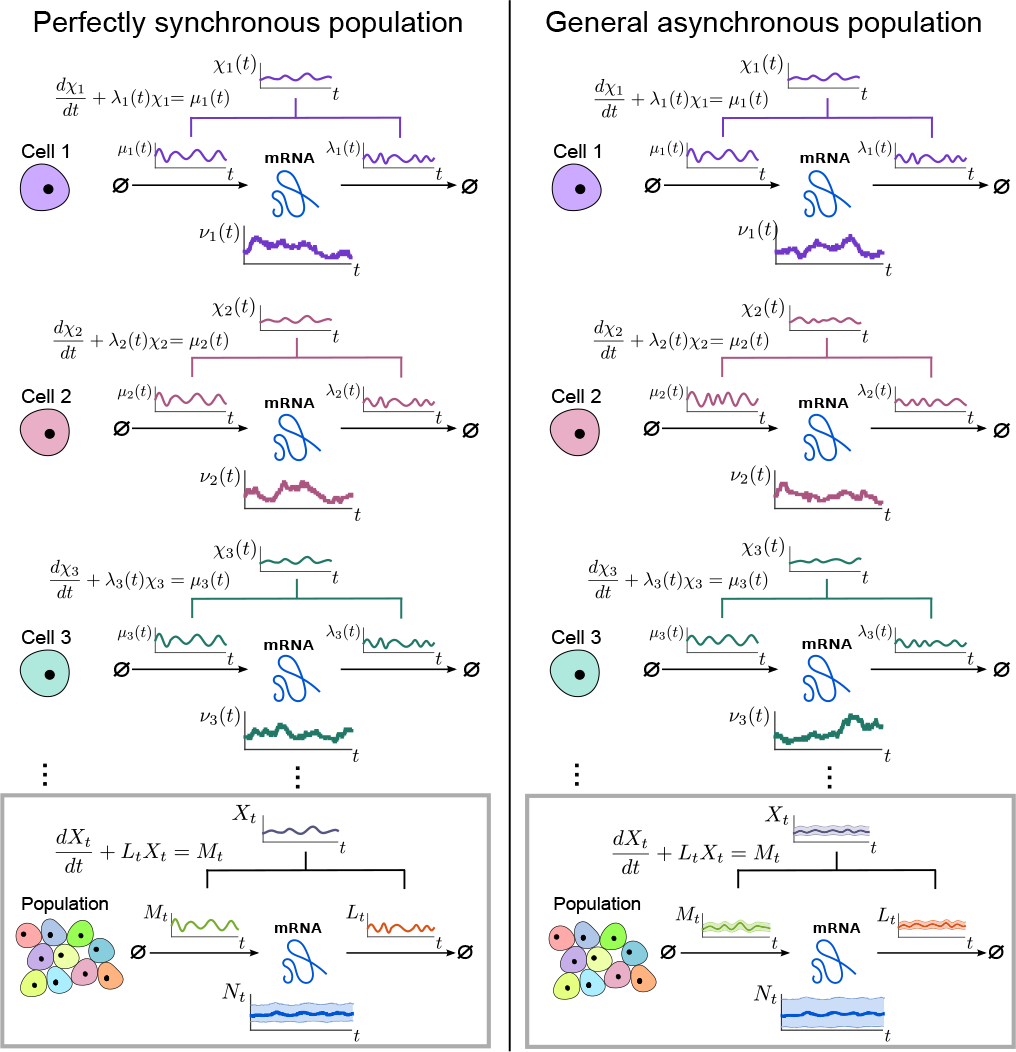
Single-cell gene transcription under upstream drives. The transcription of each cell *i* takes place under particular cellular drives {*μ_i_*(*t*)}_*t*≥0_ and {λ_*i*_(*t*)}_*t*≥0_, representing time-varying transcription and degradation rates. Both cellular drives are combined into the upstream effective drive {*χ_i_*(*t*)}>_*t*≥0_, which dictates the long-term probability distribution describing the stochastic gene expression {*v_i_*(*t*)}_*t*≥0_ within each cell (10). When there is cell-to-cell variability in the population, the cellular drives are described by processes *M* and *L* leading to the upstream effective drive *X*. The probability distribution of the population corresponds to the mixture of the upstream process *X* and the Poissonian downstream transcriptional component, as given by (14). Increased synchrony in the population implies decreased ensemble variability of the random variables *M_t_*, *L_t_*, *X_t_*, and *N_t_*.

Our aim is to obtain the probability distribution of the copy number *N_t_* under upstream time-varying cellular drives *M_t_* and *L_t_*, themselves containing stochastic parameters reflecting the cell-to-cell variability. To do this, we proceed in two steps: first, we solve the synchronous system without cell-to-cell variability; then we consider the general asynchronous case.

### A. Perfectly synchronous population

As a first step to the solution of the general case, consider a population of cells with perfectly synchronous transcription and degradation rate functions, *M* = {*μ*(*t*)}_*t*≥0_ and *L* = {λ(*t*)}_*t*≥0_; i.e., the transcription and degradation processes are defined by the same sample path for the whole population and the stochastic processes *M* and *L* have zero variance at all times (Fig. 1).

In the perfectly synchronous case, we have an immigration-death process with reaction diagram:

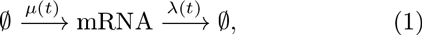

and its ME is standard:

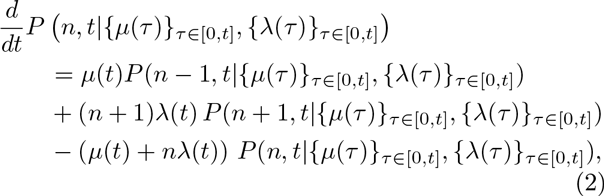

where *P*(*n*,*t*|{*μ*(*τ*)}_*τ*∈[0,*t*]_, {λ(*τ*)}_*τ*∈[0,*t*]_) denotes the probability of having *n* mRNAs at time *t* for the given history of the cellular drives {*μ*(*τ*)}_*τ*∈[0,*t*]_ and {*λ*(*τ*)}_*τ*∈[0,*t*]_.

Using the probability generating function

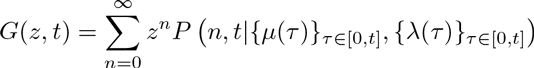

we transform the ME (2) into

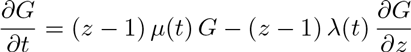

Without loss of generality, let us first consider an initial condition with *n*_0_ mRNA molecules. Using the method of characteristics, we obtain the solution:

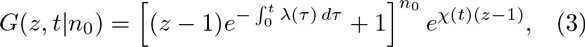

which is given in terms of

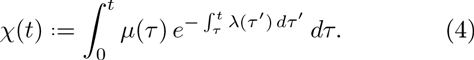

We will refer to the time-varying continuous function *χ*(*t*) as the *effective drive*, as it integrates the effect of both cellular drives.

Notice that the solution (3) can be rewritten as the product of two probability generating functions:

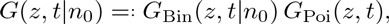

corresponding to a binomial and a Poisson distribution, respectively. Hence, for the perfectly synchronous case, the solution is given by:

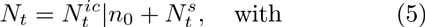

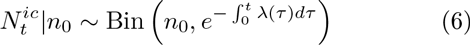

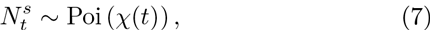

where 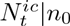 is a binomial random variable with *n*_0_ trials and success probability 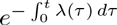, and 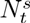 is a Poisson random variable with parameter *χ*(*t*). The physical interpretation of this breakdown is that 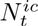 describes the mRNA transcripts that were initially present in the cell and still remain at time *t*, where as 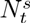 describes the number of mRNAs transcribed since *t* = 0.

Since 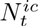 and 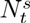 are independent, it is easy to read off the first two moments directly:

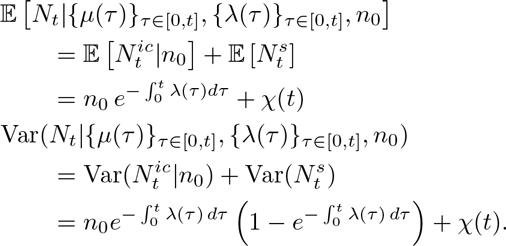

From (5)-(7), the distribution is:

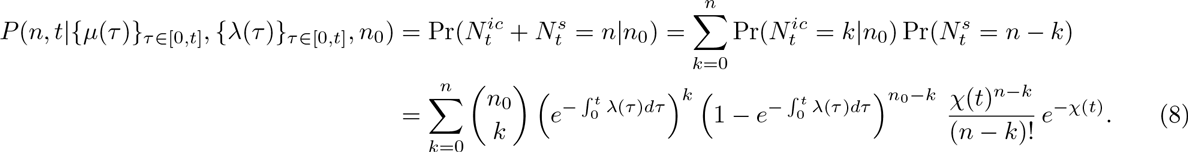

If the initial state is itself described by a random variable *N*_0_ with its own probability distribution, we apply the law of total probability to obtain the solution in full generality as (see Appendix A):

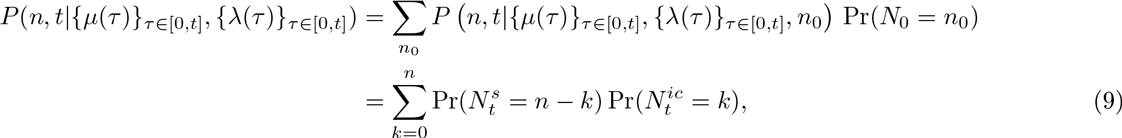

where 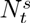 is distributed according to (5)-(7), and 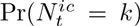 is the mixture of the time-dependent binomial distribution (5) and the distribution of the initial condition *N_0_*.

***The initial transient ‘burn in’ period*.** For biologically realistic degradation rates {λ(*t*)}_*t*≥0_, the contribution from the initial condition 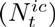 decreases exponentially. Hence as time grows the transcripts present at *t* = 0 degrade, and the population is expected to be composed of mRNAs transcribed after *t* = 0.

If the initial distribution of *N_0_* is not the stationary distribution of the ME (or, more generally, not equal to the attracting distribution of the ME, as defined in Appendix A), there is an initial time-dependence of *P*(*n, t*) lasting over a time scale *T^ic^* (given by 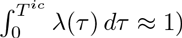, which corresponds to a ‘burn-in’ transient associated with the decay of the initial condition. For instance, the time-dependence described in Ref. [13] for the random telegraph model (Fig. 6) corresponds to this ‘burn-in’ transient.

On the other hand, when the initial distribution of *N*_0_ is the stationary distribution (or the attracting distribution) of the ME, the component containing the initial condition 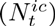 and the long-term component 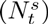 balance each other at every point in time, maintaining stationarity (or the attracting distribution), as shown analytically in Appendix A.

***The long-term behaviour of the solution*.** In this work, we focus on the time dependence of *P*(*n*, *t*) induced through non-stationarity of the parameters and/or correlated behaviour of the cells within the population. Hence for the remainder of the paper, we neglect the transient terms. Consequently, for perfectly synchronous cellular drives, the solution of the ME (2) is a Poisson random variable with time-dependent rate equal to the effective upstream drive, *χ*(*t*):

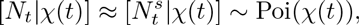

with distribution

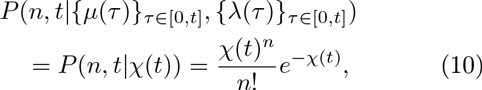

which makes explicit the dependence on the *history* of the sample paths {*μ*(*τ*)}_*τ*∈[0,*t*]_, {λ(*τ*)}_τ∈[0,*t*]_, which is encapsulated in the *value* of the effective drive *χ*(*t*) at time *t*.

Indeed, from (4) it follows that the sample path {*χ*(*t*)}_*t*≥0_ satisfies a first order linear ordinary differential equation with time-varying coefficients:

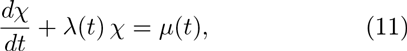

which is the rate law for a chemical reaction with zeroth-order production with rate *μ*(*t*), and first-order degradation with rate λ(*t*) per mRNA molecule. For biologically realistic (i.e., positive and finite) cellular drives, *χ*(*t*) is a continuous function.

### B. The general asynchronous case:cell-to-cell variability in the cellular drives

Consider now the general case where different sample paths for the cellular drives are possible, i.e., the cell population has some degree of asynchrony and *M_t_* and *L_t_* have non-zero variance for at least some *t* ≥ 0. The transcription and degradation rates are then described by stochastic processes *M* and *L*:

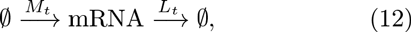

and the collection of all differential equations of the form (11) is represented formally by the random differential equation^1^

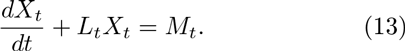

Equations of this form appear in many sciences, and a large body of classical results allows us to determine the probability density function of the upstream process, *X_t_* [37–39]. Below, we use such results to obtain *f_X_t__*(*x, t*) for biologically relevant models.

Note that from Eq. (10) and the law of total probability, it follows that the probability mass function for the random variable *N_t_* under cellular drives described by the random processes *M* and *L* is given by the *Poisson mixture* (or compound) distribution:

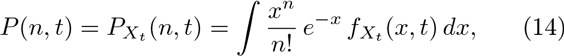

where the density *f_X_t__* (*x,t*) of the effective drive *X_t_* (to be determined) can be understood as a *mixing density*. The notation *P_X_t__* (*n*, *t*) recalls explicitly the dependence of the solution on the density of *X_t_*, but we drop this reference and use *P*(*n, t*) below to simplify notation. The problem of solving the full ME is thus reduced to finding the mixing density *f_X_t__* (*x,t*). Note that for synchronous drives, we have *f_X_t__*(*x, t*) = *δ*(*χ*(*t*)−*x*), where *δ* is the Dirac delta function, and (14) reduces to (10).

Equation (14) also shows that there are two separate sources of variability in gene expression, contributing to the distribution of *N_t_*. One source of variability is the Poisson nature of transcription and degradation, common to every model of the form considered here; the second source is the time-variation or uncertainty in the cellular drives, encapsulated in the upstream process *X_t_* describing the ‘degree of synchrony’ between cells and/or their variability over time. In this sense, Eq. (14) connects with the concept of separable ‘intrinsic’ and ‘extrinsic’ components of gene expression noise pioneered by Swain *et al.* [40–43]. Yet rather than considering moments, the full distribution *P*(*n,t*) is separable into a model-dependent ‘upstream component’ given by *f_X_t__*(*x,t*), and a downstream transcriptional ‘Poisson component’ common to all models of this type.

## III. THE EFFECTIVE UPSTREAM DRIVE IN GENE TRANSCRIPTION MODELS

The generic model of gene transcription and degradation with time-dependent drives introduced above provides a unifying framework for several models previously considered in isolation. In this section, we exemplify the tools to obtain the density of the effective drive *f_X_t__* (*x, t*) analytically or numerically through relevant examples.

### A. Gene transcription under upstream drives with static randomness

In this first section, we consider models of gene transcription where the upstream drives are deterministic, yet with random parameters representing cell variability

#### 1. *Random entrainment to upstream sinusoidal drives: random phase offset in transcription or degradation rates*

Equation (13) can sometimes be solved directly to obtain *f_X_t__* (*x, t*) from a transformation of the random variables *M_t_* and *L_t_*. We now show two such examples, where we explore the effect of entrainment of gene transcription and degradation to an upstream periodic drive [44].

First, consider a model of gene transcription of the form (12) with transcription rate given by a sinusoidal function and where each cell has a random phase. This *random entrainment* (RE) model is a simple representation of a cell population with transcription entrained to an upstream rhythmic signal, yet with a random phase offset for each cell:

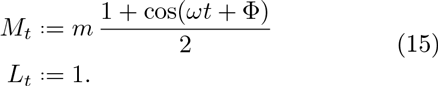

Here *m* and *ω* are given constants and Φ is a (static) random variable describing cell-to-cell variability (or uncertainty). Solving Eq. (13) in this case, we obtain

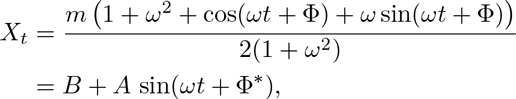

where 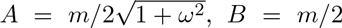 and Ф* = Ф + arctan(1/*ω*).

Suppose Ф* is uniformly distributed on [−*r, r*], *r* ≤ *π*. Inverting the sine with Ф* restricted to [−*r, r*], we obtain

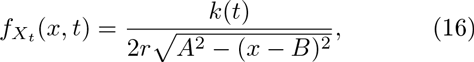

where *k*(*t*) ∈ {0,1,2} is the number of solutions of sin *θ* = (*x* − *B*)/*A* for *θ* ∈ (*ωt* − *r*, *ωt* + *r*). As the phase distribution of the drives becomes narrower, the upstream variability disappears: *r* → 0 ⟹ *f_X_t__* (*x, t*) → *δ*((*B* + *A* sin *ωt*) − *x*). In this limit, all cells follow the entraining drive exactly, and *P*(*n, t*) becomes a Poisson distribution at all times

Figure 2 depicts *f_X_t__* for *r* = 0 (no cell-to-cell phase variation, (a)) and for *r* = *π*/2, and *r* = *π* (increasingly wider uniform distribution of phases, (b) and (c)). The full distribution *P*(*n, t*) is obtained using (16) and (14).

Second, let us consider the same model of entrainment to an upstream sinusoidal signal with a random offset, but this time via the degradation rate:

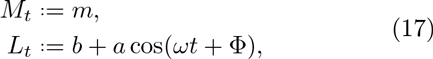

with *m*, *a*, *b*, and *ω* given constants, and Φ a (static) random variable.

Eq. (13) can be solved approximately [44] to give

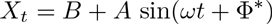

where 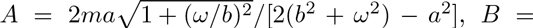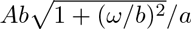 and Ф* = *π*/2 − arg[cos(*π*/2 − arctan(*b*/*ω*) − Ф)]. As before, if Ф* is uniform on [−*r,r*], *r* ≤ *π*, the density of the effective drive takes the same form (16) as above.

#### 2. *Upstream Kuramoto promoters with varying degree of synchronization*

As an illustrative computational example, we study a population of cells whose promoter strengths display a degree of synchronization across the population. To model this upstream synchronization, consider the *Kuramoto promoter model*, where the promoter strength of each cell *i* depends on an oscillatory phase *θ*_*i*_(*t*), and cells are coupled via a Kuramoto model [45–47]. We then have a model of the form (12) with:

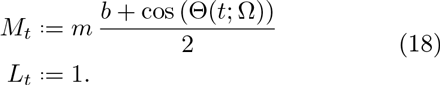

Here m; b are constants and 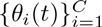 are the phase variables for the *C* cells governed by the globally coupled Kuramoto model:

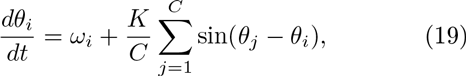

where *K* is the coupling parameter and the intrinsic frequency of each cell, *ω_i_*, is drawn from the random distribution 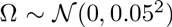. The Kuramoto model allows us to tune the degree of synchrony through the coupling *K*: for small *K*, the cells do not display synchrony since they all have a slightly different intrinsic frequency; as *K* is increased, the population becomes more synchronized.

**FIG. 2.**
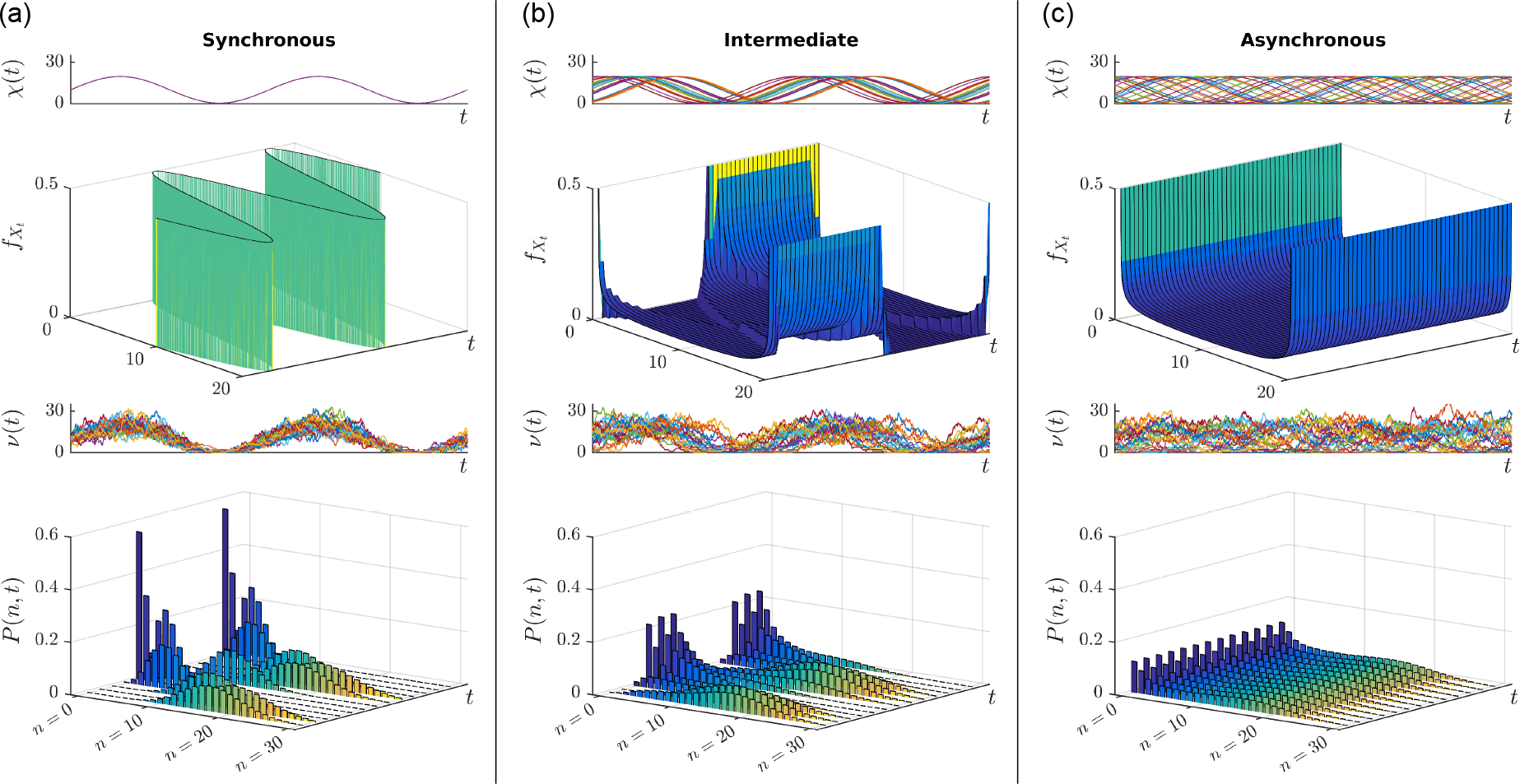
Gene transcription under the random entrainment model (15) with constant degradation rate and transcription rates entrained to an upstream sinusoidal signal with *ω* = 1. Each cell has a random phase offset *ϕ* drawn from a distribution. (a) The synchronous population corresponds to identical phases across the population. In this case, the transcription reflects the time variability of the upstream drive mixed with the stochasticity due to the downstream Poisson process of transcription. When the random phases *ϕ* are uniformly distributed on an interval of range (b) *π* and (c) 2*π*, the population becomes increasingly asynchronous. For all three cases, we show (top to bottom): sample paths of the effective drive, *X*; its density *f_X_t__*(*x,t*) given by Eq. (16) sample paths of the number of mRNAs, *N*; and the full solution of the ME *P*(*n,t*).

This model is a simple representation of nonlinear synchronization processes in cell populations with intrinsic heterogeneity [48–51]. In Figure 5(a), we show how the sample paths change as the degree of synchrony increases, and we exemplify the use of (14) for the numerical solution of the gene expression of this model

### B. Asynchronous transcription under stochastic multistate promoters

In the previous section, we obtained *f_X_t__* (*x, t*) by capitalizing on the precise knowledge of the sample paths of *M* and *L* to solve (13) explicitly. In other cases, we can obtain *f_X_t__* (*x, t*) by following the usual procedure of writing down an evolution equation for the probability density of an *expanded* state that is Markovian, and then marginalizing. More specifically, let the vector process **Y** prescribe the upstream drives, so that *M* = *M*(**Y**, *t*) and *L* = *L*(**Y**, *t*), and consider the expanded state **X**_*t*_: = (*X_t_*,**Y**_*t*_). Note that since **Y** is upstream, it prescribes *X* (and not vice versa). We can then write the evolution equation for the joint probability density *f*_X_t__(*x, y, t*):

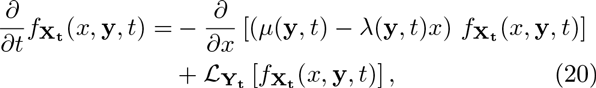

which follows from conservation of probability. In Equation (20), the differential operator for *X*, which follows from (13), is the first jump moment [52] conditional upon **Y**_t_ = y (and hence upon *M_t_* = *μ*(*y, t*) and *L_t_* = λ(*y, t*)); the second term 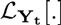 is the infinitesimal generator of the upstream processes. In particular, for continuous stochastic processes 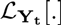 is of Fokker-Planck type, and for Markov chains 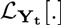 is a transition rate matrix. The desired density *f_X_t__* (*x, t*) can then be obtained via marginalization.

Equation (20) can be employed to analyze the widely used class of transcription models with asynchronous, random promoter switching between discrete states, where each state has different transcription and degradation rates representing different levels of promoter activity due to, e.g., transcription factor binding or chromatin remodeling [34]. A classic example is the *random telegraph* (RT) model, first used by Ko in 1991 [53] to explain cell-to-cell heterogeneity and bursty transcription (Fig. 3a).

In our framework, such random promoter switching can be understood as an upstream *stochastic* process driving transcription as follows. Let us assume that the promoter can attain *D* states *s*, and each state has constant transcription rate *m_s_* and constant degradation rate *ℓ_s_*. The state of the promoter is described by a random process *S* = {*S_t_* ∈ {1, 2,…,*D*}: *t* ≥ 0}, with sample paths denoted by {*ς*(*t*)}_*t*≥0_ and its evolution is governed by the *D*-state Markov chain with transition rate *k_sr_* from state *r* to state *s*. The state of the promoter *S_t_* = *s* prescribes that *M_t_* = *m_s_* and *L_t_* = *ℓ_s_*. Hence, the sample paths of *M* and *L* are a succession of step functions with heights *m_s_* and *ℓ_s_*, respectively, occurring at exponentially distributed random times.

As described above, we expand the state space of the cellular drives to include the promoter state **X**_*t*_ = {*X_t_, S_t_*}. The evolution equation (20) is then given by *D* coupled equations:

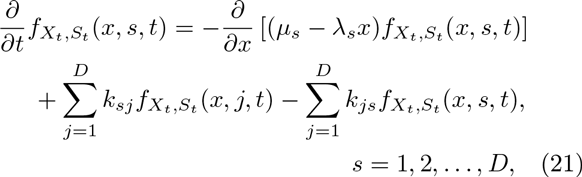

which can be thought of as a set of multistate Fokker-Planck-Kolmogorov equations [52]. Marginalization then leads to the density of the effective drive:

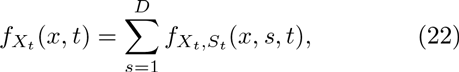

and the full ME solution is obtained from (22) and (14).

We illustrate this approach more explicitly with two examples (Fig. 3): a re-derivation of the known solution of the standard RT model; and the solution of the 3-state cyclic model with a refractory state. Results for other promoter architectures are discussed in [54].

#### 1. *The Random Telegraph model (2 states)*

Although the RT model has been solved by several methods [13, 32, 33], we briefly rederive its solution within the above framework to clarify its generalisation to other promoter architectures.

Consider the standard RT model (Fig. 3a), with promoter switching stochastically between the active state *s*_on_ = 1, with constant transcription rate *m_1_* = *m*, and the inactive state *s*_off_ = 0, where no transcription takes place, *m*_0_ = 0. The transition rates between the two states are *k*_10_ = *k*_on_ and *k*_01_ = *k*_off_. Without loss of generality, we assume *ℓ*_1_ = *ℓ*_0_ = λ(*t*) ≡ 1;. The transcription model is of the form (12) with:

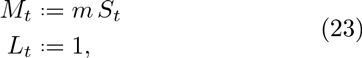

where *S* = {*S*_t_ ∈ {0,1}: *t* ≥ 0} with waiting times drawn from exponential distributions: *τ*_off_ ~ exp(1/*k*_on_) and *τ*_on_ ~ exp(1/*k*_off_)

**FIG. 3.**
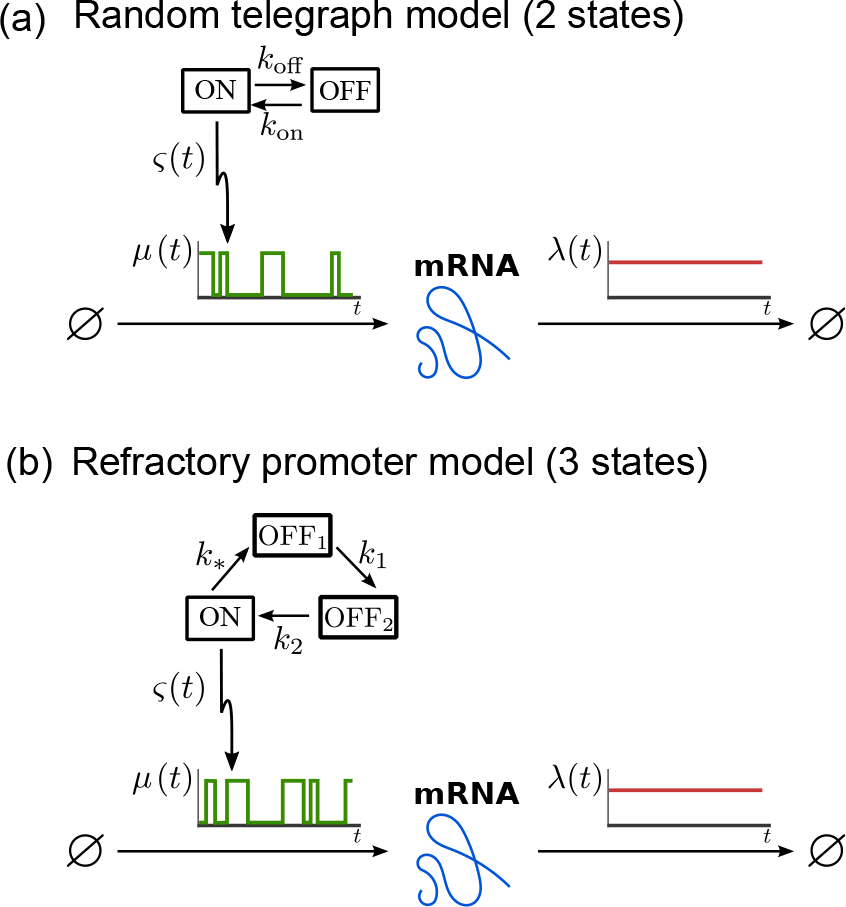
Asynchronous stochastic promoter switching models correspond to upstream stochastic processes. The promoter cycles between the discrete states, transitioning stochastically with rates as indicated: (a) the standard 2-state random telegraph (RT) model; (b) the 3-state refractory promoter model.

Let *Z_t_* = *X_t_/m*, and denote *f*_on_(*z,t*): = *f_Z_t_,S_t__*(*z,s*_on_,*t*) and *f*_off_(*z,t*): = *f_Z_t_,S_t__*(*z,s*_off_,*t*), with *z* ∈ (0,1). Then the multistate Fokker-Planck-Kolmogorov equations (21) are:

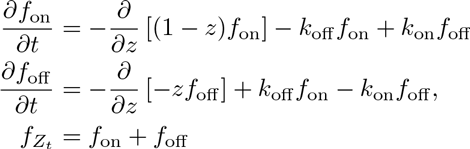

with integral conditions 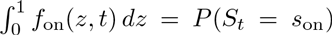 and 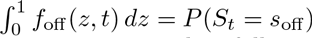.

At stationarity, it then follows [55] that

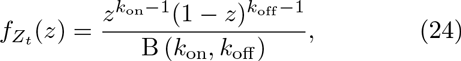

where B(*a,b*) = Γ(*a*)Γ(*b*)/Γ(*a* + *b*) is the Beta function. In other words, at stationarity, the normalised effective drive is described by a Beta distribution:

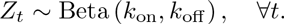

Using (24) and (14), we obtain that the full stationary solution is the Poisson-Beta mixture:

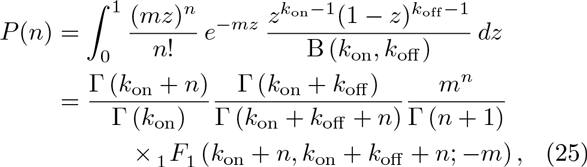

where _1_*F*_1_ (*a, b; z*) is the confluent hypergeometric function [56].

#### 2. *The Refractory Promoter model (3 states)*

In the standard RT model, the waiting times in each state are exponentially distributed. In recent years, time course data have shown that the *τ*_off_ do not conform to an exponential distribution, leading some authors to incorporate a second inactive (refractory) state, which needs to be cycled through before returning to the active state [36, 57]. The net ‘OFF’ time is then the sum of two exponentially distributed waiting times.

In this *refractory promoter model* (Fig. 3b), the promoter switches through the states *s*_*_, *s*_1_, and *s*_2_ with rates *k*_*_, *k*_1_ and *k*_2_. Transcription takes place at constant rate *m* only when the promoter is in the active state *s*_*_ and, without loss of generality, we assume a constant degradation rate λ(*t*) ≡ 1 for all states. This model is of the same form as (23), and is solved similarly.

Making the change of variables *Z_t_* = λ*X_t_/m* = *X_t_/m* and, using the notation *f_i_*(*z, t*): = *f_Z_t__,_S_t__* (*z, t, s_i_*), the multistate Fokker-Planck-Kolmogorov equations are

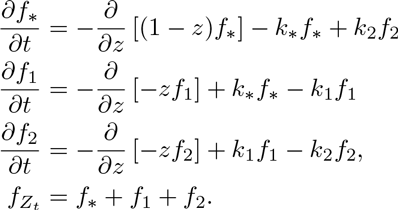

with three integral conditions 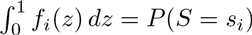.

At stationarity, we find

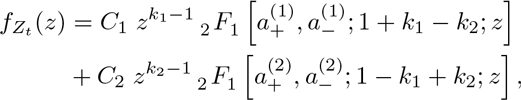

with 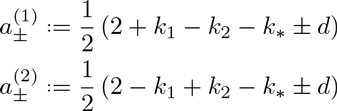, where 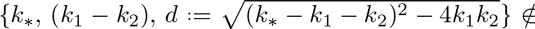 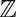 and _2_*F*_1_ (*a, b*; *c*; *z*) is the Gauss hypergeometric function [56]. The full stationary solution *P*(*n*) is then obtained from (14).

For a detailed derivation (including expressions for the integration constants *C*_1_ and *C*_2_), see Appendix B.

### C. Asynchronous multistate models with upstream promoter modulation

Finally, we consider a model of gene transcription that incorporates features of models described in Sections IIIA and III B. Such a situation is of biological interest and appears when individual cells exhibit correlated dynamics in response to upstream factors (e.g., changing environmental conditions, drives or stimulations), but still maintain asynchrony in internal processes, such as transcription factor binding [31, 58].

**FIG. 4.**
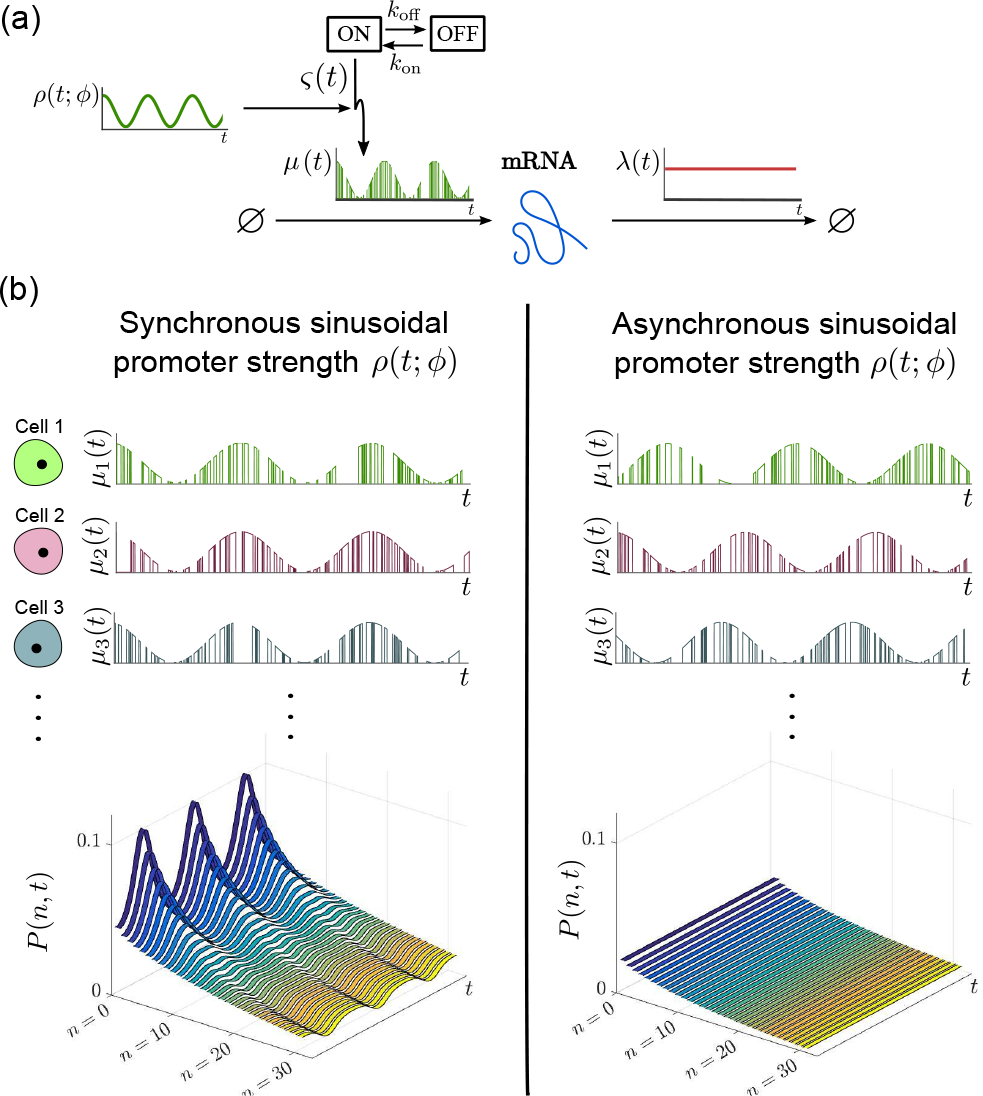
(a) Modulated Random Telegraph (MRT) model: each cell switches asynchronously between ‘ON’ and ‘OFF’ states, but the magnitude of the ‘ON’ transcription rate is modulated by the function *ρ*(*t; ϕ*), a sinusoid representing an upstream periodic process. The phase *ϕ* represents the cell-to-cell variability and leads to the varying degree of synchrony across the population. (b) Sample paths *μ_i_*(*t*) and solution of the probability distribution *P*(*n, t*) of the MRT model for synchronous (left) and asynchronous (right) modulation. In the asynchronous case, the upstream drive has a random phase across the cells with distribution 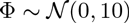.

To illustrate this concept, we consider the *modulated random telegraph* (MRT) model, a combination of the RE model (15) and the RT model (23), i.e., the promoter strength is modulated by an upstream sinusoidal drive with random phase Φ, as in the RE model, yet the promoter switches stochastically between active/inactive states with rates *k*_on_ and *k*_off_, as in the RT model. In this model, the transcription rate is correlated between cells through the entrainment to the upstream sinusoidal drive as a simple model for, e.g.,circadian gene expression:

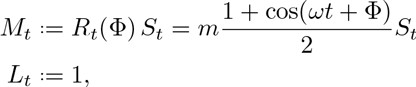

where *m,ω* > 0 are constants; Φ is the random phase across the cell population; and *S* = {*S_t_* ∈ {0,1}: *t* ≥ 0}, with exponential waiting times, describes the stochastic promoter switching (Fig. 4a).

The solution of this model follows from the RT probability density (24) conditioned on the random phase Φ, which prescribes the sample path {*ρ*(*t*,*ϕ*)}_*t*≥0_ of the promoter strength *R*. The resulting scaled Beta distribution

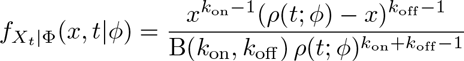

is then marginalized over the phase Φ to obtain the density *f_X_t__* (*x,t*) of the effective drive. For instance, if the phases are normally distributed 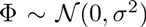, we have (see Fig. 4b):

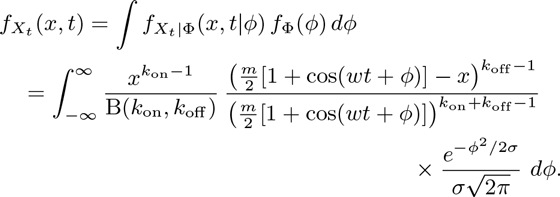

As *σ* → ∞, the population becomes asynchronous in the promoter strength, as well as in the state transitions, and time dependence wanes(Fig. 4b).

## IV. ENSEMBLE NOISE CHARACTERISTICS IN TIME-VARYING POPULATIONS

In the previous sections, we were concerned with the full time-dependent probability distribution *P*(*n, t*) for the mRNA copy number *N*. However, in many circumstances such detailed information is not required, and simpler characterizations based on ensemble averages (e.g., Fano factor, coefficient of variation) are of interest. Simple corollaries from the Poisson mixture expression (14) allow us to derive expressions for the ensemble moments and other noise characteristics, as shown below. We remark that, in this section, all the expectations are taken over the distribution describing the cell population.

### A. Time-dependent ensemble moments over the distribution of cells

To quantify noise characteristics of gene expression in a population, the ensemble moments 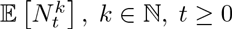 are often determined via the probability generating function [14, 32, 59] or by integrating the master equation [19, 34, 60]. However, stationarity is usually assumed and the moments derived are not suitable for time-varying systems. Here we use corollaries of the Poisson mixture expression (14) to derive expressions for the ensemble moments for time-varying systems under upstream drives.

From (10) we have [*N_t_*|*X_t_* = *x*] ~ Poi(*x*); hence

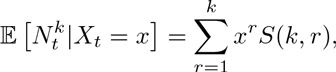

where 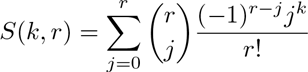 are the Stirling numbers of the second kind [61]. The law of total probability then gives

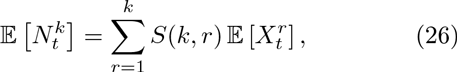

or, equivalently, 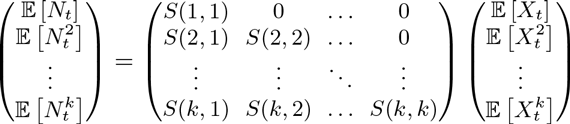.

Therefore the ensemble moments of the mRNA copy number 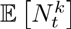 can be obtained in terms of the moments of the effective drive 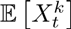, and vice versa. For instance, 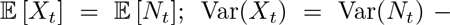 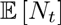 and the skewness 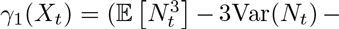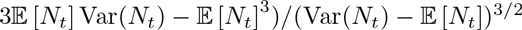.

The expressions (26) complement those used by Swain *et al.* in their seminal work on intrinsic/extrinsic noise [41], and those by Hilfinger and Paulsson to separate intrinsic/extrinsic fluctuations [42].

### B. Time-dependent ensemble Fano factor: a measure of synchrony in the population

A commonly used measure of variability in the population is the *ensemble Fano factor*:

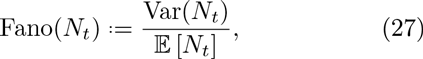

which is unity for the Poisson distribution. Its use has been popularized as a measure of the deviation from the stationary solution of the transcription of an unregulated gene with constant rates [62, 63], which is Poisson; hence with Fano(*N_t_*) ≡ 1, ∀*t*.

For time-varying systems, however, the ensemble Fano factor conveys how the dynamic variability in single cells combines at the population level. Indeed, Fano(*N_t_*) can be thought of as a measure of synchrony in the population at time *t*. For instance, it follows from Eq. (10) that the ensemble Fano factor for a population with perfectly synchronous drives is always equal to one, Fano(*N_t_*) ≡ 1, ∀*t*. Even if the upstream drive *χ*(*t*) changes in time, the population remains synchronous and has a Poisson distribution at all times. On the other hand, under the assumptions of our model, when Fano(*N_t_*) varies in time it reflects a change in the degree of synchrony between cells, as captured by the upstream drive *X_t_*. Indeed, from (26) it follows that:

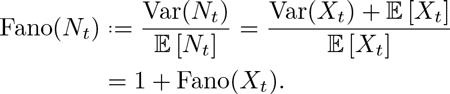

**FIG. 5.**
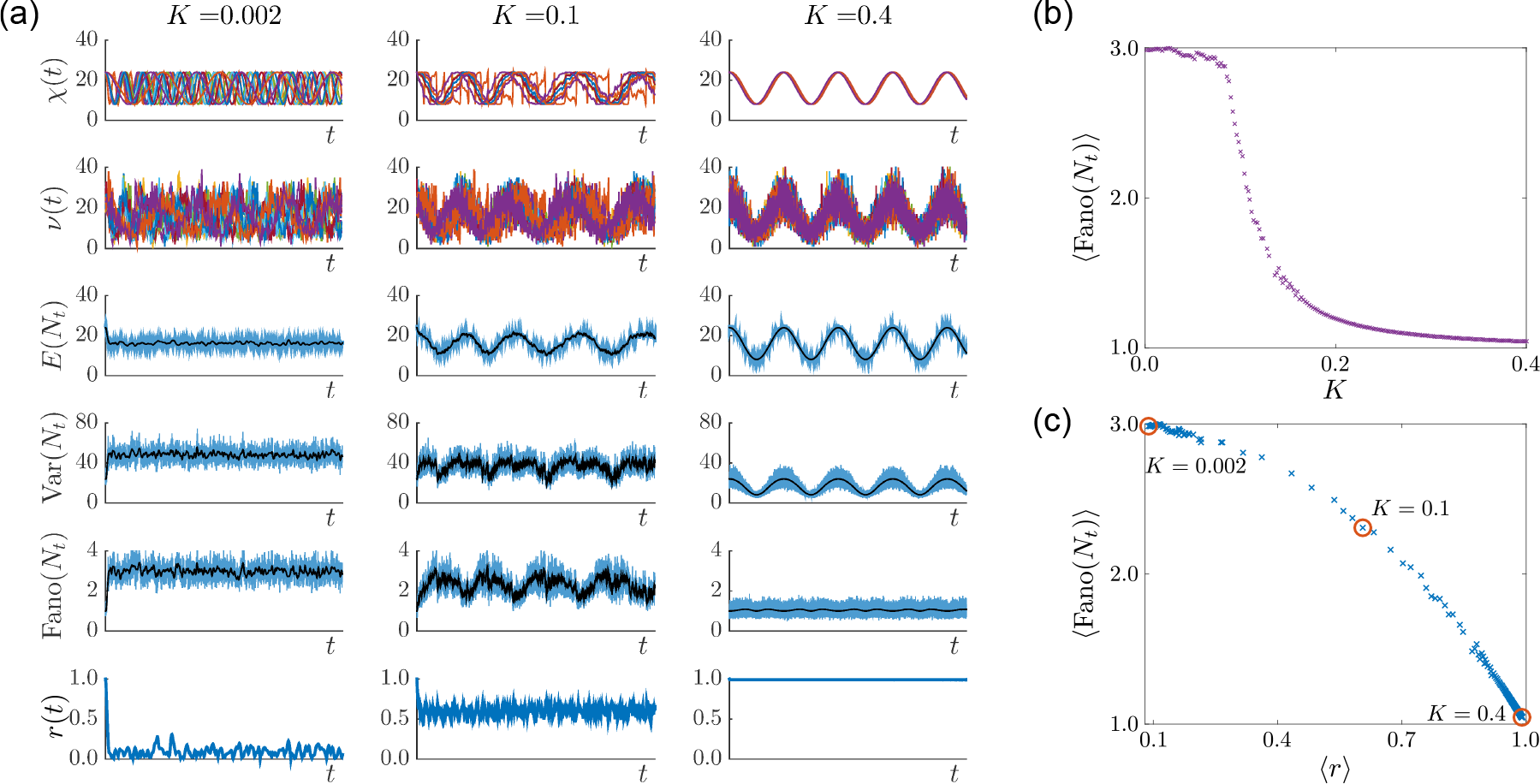
Noise characteristics of the Kuramoto promoter model (18). (a) Numerical simulations for *C* = 100 oscillatory cells and different coupling parameters: *K* = 0:002,0.1,0.4 (left, middle, right columns). For each coupling, the sample paths of the upstream effective drive *X* and mRNA counts *N* are shown. The mean, variance, and ensemble Fano factor of *N* were calculated from the sample paths of *N* (blue lines) and, more efficiently, from the sample paths of *X* (black lines). The last row shows the Kuramoto order parameter *r*(*t*) measuring the cell synchrony, signaled by *r*(*t*) → 1. (b) Ensemble Fano factor (averaged over the simulated time courses) against coupling parameter *K* ∈ (0, 0.4]. As *K* is increased, the oscillators become synchronized and the ensemble Fano factor decreases towards the Poisson value of unity. (c) Scatter plot of the ensemble Fano factor against the order parameter *r*(*t*) (both averaged over the simulated time courses). As the oscillators become synchronized 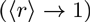, the ensemble Fano factor also approaches 1, signifying that the distribution is Poissonian at all times.

The greater the synchrony at time *t*, the closer Fano(*N_t_*) is to unity, since the deviation from the Poisson distribution emanates from the ensemble Fano factor of the upstream drive *X_t_*.

As an example, consider the Kuramoto promoter model (18)-(19) introduced in Section IIIA 2, where the cells in the population become more synchronized as the value of the coupling *K* is increased. Figure 5 shows simulation results for 100 cells with a range of couplings. The order parameter *r*(*t*) ∈ [0,1] measures the phase coherence of the oscillators at time *t*; as it grows closer to 1, so grows the degree of synchrony. Using the Kuramoto numerics, we calculate the ensemble Fano factor Fano(*N_t_*) for the transcription model. As seen in Fig. 5(b)-(c), the more synchronous, the closer the Fano factor is to the Poisson value: 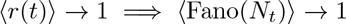.

Figure 5 also illustrates the computational advantages of our method. The cost to approximate the time-varying ensemble moments is drastically reduced by using (26), because transcription and degradation events do not have to be simulated. The sample paths of the effective drive *χ_i_*(*t*) were used to estimate the time-varying moments: 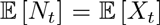 and 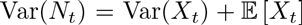 (shown in black). These correspond to the numerical simulation of ODEs, and are far more efficient than sampling from realisations *v_i_*(*t*) of the mRNA copy number.

### V. VARIABILITY OVER TIME: STATIONARITY AND ERGODICITY

Our results up to now have not assumed stationarity— in general the distribution (14) and ensemble moments (26) depend on time. If the cells in the population are uncorrelated and *M* and *L* are *stationary* (i.e., their statistics do not change over time), then *f_X_t__* (*x,t*) tends to a stationary density *f_X_t__* (*x*) [38], and the full solution *P*(*n,t*) also tends to a stationary distribution *P*(*n*).

Under such assumptions leading to stationarity, any time dependence in the solution *P*(*n,t*) describes the ‘burn-in’ transient from an initial condition towards the attracting stationary distribution, as discussed in Section II A. An example of such transience in the random telegraph model (23) appeared in [13] describing how the distribution *P*(*n,t*) settles from an initial Kronecker delta distribution *P*(*n*, 0) = *δ*_*n*0_ to the stationary distribution (25) (see Fig. 6 and Appendix A 2).

If, in addition to stationarity, we assume the cells to be indistinguishable, the population is *ergodic.* In this case, the distribution obtained from a single cell over a time *T*, as *T* → ∞, is equivalent to the distribution obtained from a time snapshot at stationarity of a population of *C* cells, as *C* → ∞:

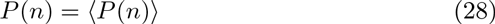

where 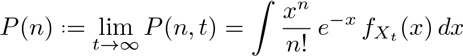
and

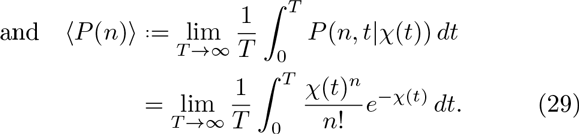

where 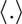 denotes time-averaging. Under the assumption of ergodicity, the averages computed over single-cell sample paths can be used to estimate the stationary distribution of the population.

**FIG. 6.**
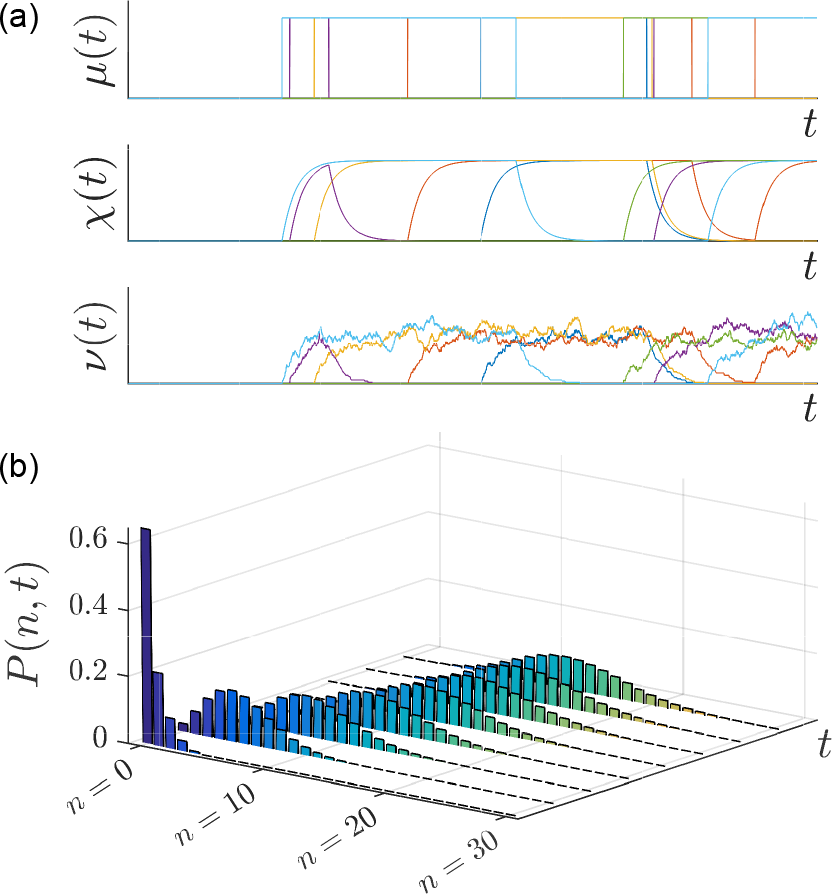
‘Burn-in’ transient in the random telegraph (RT) model. (a) Sample paths of the transcription rate *M*, the effective upstream drive *X*, and the number of mRNAs *N* for an initial condition *P*(0,0) = 1 with all cells initialised in the inactive state [13]. (b) The full solution of the RT model for this initial probability distribution exhibits an exponential decay as the system approaches its stationary distribution. The delta distribution at *t* = 0 is omitted for scaling purposes

### A. Ergodic systems: stochastic *vs* deterministic drives

It has been suggested that stochastic and periodic drives lead to distinct properties in the noise characteristics within a cell population [42]. We investigate the effect of different temporal drives on the full distribution (14) under ergodicity using (28)-(29). Note that when *χ*(*t*) is periodic with period *T*, the limit in Eq. (29) is not required. In Figure 7, we show the time-averaged distribution 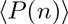 for a cell under three different upstream drives *μ*(*t*): (i) a continuous sinusoidal form; (ii) a discontinuous square wave form; (iii) a random telegraph (RT) form, which can be thought of as the stochastic analogue of the square wave. In all cases, the drive {*μ*(*t*)}_*t*≥0_ ∈ [0,20] with the same period, or expected period, *T*. For simplicity, we set λ(*t*) ≡ 1.

**FIG. 7.**
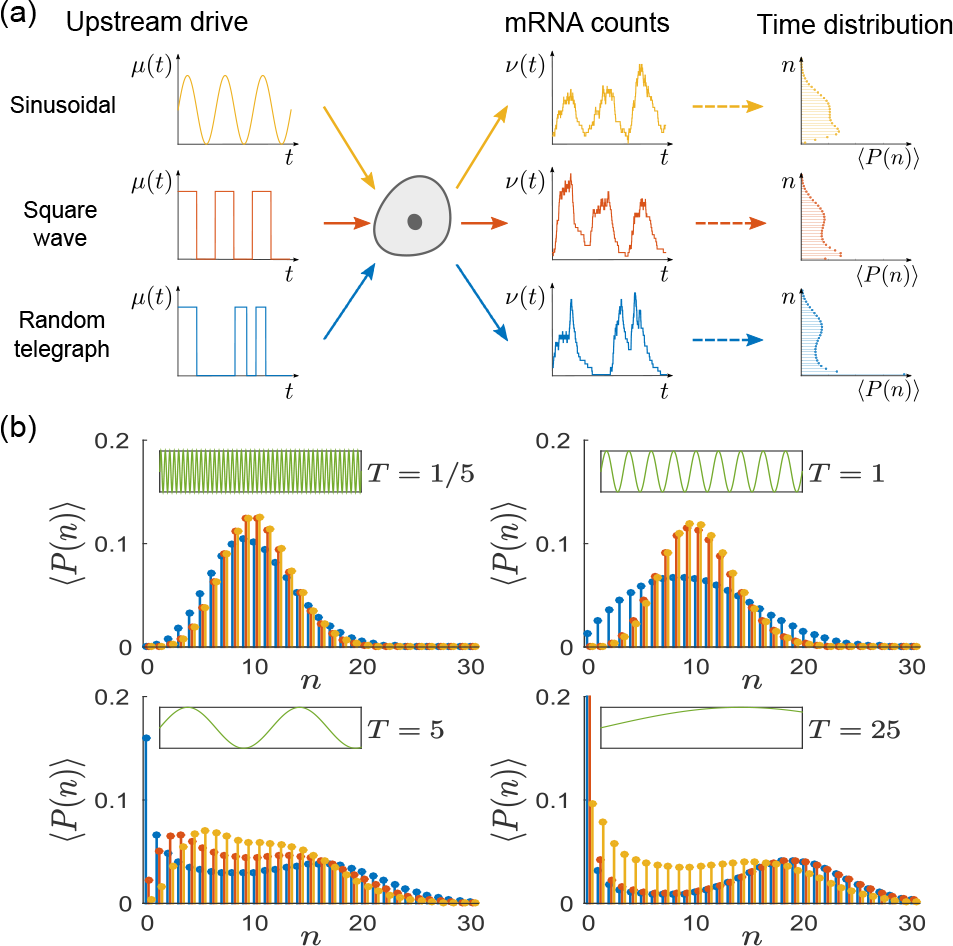
Ergodic transcription models under periodic and stochastic upstream drives. (a) We consider gene transcription under three drives *μ*(*t*) ∈ [0,20]: a sinusoidal wave with period *T* (yellow); a square wave with period *T* (red); a random telegraph process with expected waiting time *T*/2 in each state (blue). For such ergodic systems, the distribution computed over time 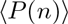 corresponds to the stationary distribution. (b) The distribution 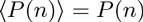 presents distinct features as the period *T* is varied.

As the period *T* is varied, the similarity between the distributions under the three upstream drives varies considerably (Fig. 7). At small *T*, the distributions under sinusoidal and square wave forms are most similar; whereas at large *T*, the distributions under square wave and RT forms become most similar. In general, the distribution of the RT model has longer tails (i.e., *n* low and high) as a consequence of long (random) waiting times that allow the system to reach equilibrium in the active and inactive states, although this effect is less pronounced when the promoter switching is fast relative to the time scales of transcription and degradation (e.g., *T* = 1/5). On the other hand, as *T* grows, the square wave and RT drives are slow and the system is able to reach the equilibrium in both active and inactive states. Hence the probability distributions of the square wave and RT drives become similar, with a more prominent bimodality.

### B. The temporal Fano factor: windows of stationarity in single-cell time course data

The *temporal Fano factor* (TFF) is defined similarly to the ensemble version (27), but is calculated from the variance and mean of a *single time series* {*v* (*t*)}_t_)_*t*≥0_;over a time window *W*: = (*t*_1_, *t*_2_):

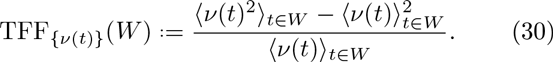

In fact, this is the original definition of the Fano factor [64], which is used in signal processing to estimate statistical fluctuations of a count variable over a time window. Although *N_t_* is not a count variable (it decreases with degradation events), the TFF can be used to detect windows of stationarity in single-cell time courses.

Figure 8a shows a single-cell sample path {*v*(*t*)}_*t*≥0_ generated by the (leaky) random telegraph model with constant degradation rate λ, and transcription rates *μ*_1_ > *μ*_0_ > 0 for the active and inactive promoter states. The leaky RT model is equivalent to the standard RT model, and switches between two states with expectations *μ*_1_/λ and *μ*_0_/λ. In the time windows *W* between promoter switching, {*v*(*t*)}_*t*∈*W*_ can be considered almost at stationarity and described by a Poisson distribution with parameter *μ*_0_/λ (resp. *μ*_1_/λ) in the inactive (resp. active) state. Hence TFF_{*ν*(*t*)}_ (*W*) ≃ 1 across most of the sample path, except over the short transients *W*_trans_ when the system is switching between states, where TFF_{*v*(*t*)}_(*W_trans_*) > 1 (Fig. 8b).

Alternatively, this information can be extracted robustly from the *cumulative Fano factor* (cTFF):

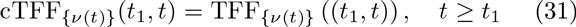

which is computed over a *growing window* from a fixed starting time *t*_1_. The cTFF is a cumulative moving average giving an integrated description of how the stationary regimes are attained between switching events indicated by the step-like structure of the heatmap in Fig. 8c.

## VI. DISCUSSION

In this paper, we have presented the solution of the master equation for gene transcription with upstream dynamical variability, in a setting that allows a unified treatment of a broad class of models. The framework allows quantitative biologists to go beyond stationary solutions when using inference to analyze noise in singlecell experiments. As an alternative to computational approaches where many cells are explicitly simulated to account for observed variability, our work takes a parsimonious approach and uses a simple gene transcription model of Poissonian type, but considers explicitly the effect of dynamical and cell-to-cell upstream variability in the solution of the master equation. The solution obtained can be viewed as combining an upstream component (dynamic or static, deterministic or stochastic) with a downstream Poissonian immigration-death process. This structure can describe both time-dependent snapshots of the population as well as the variability over single-cell time courses in a coherent fashion.

**FIG. 8.**
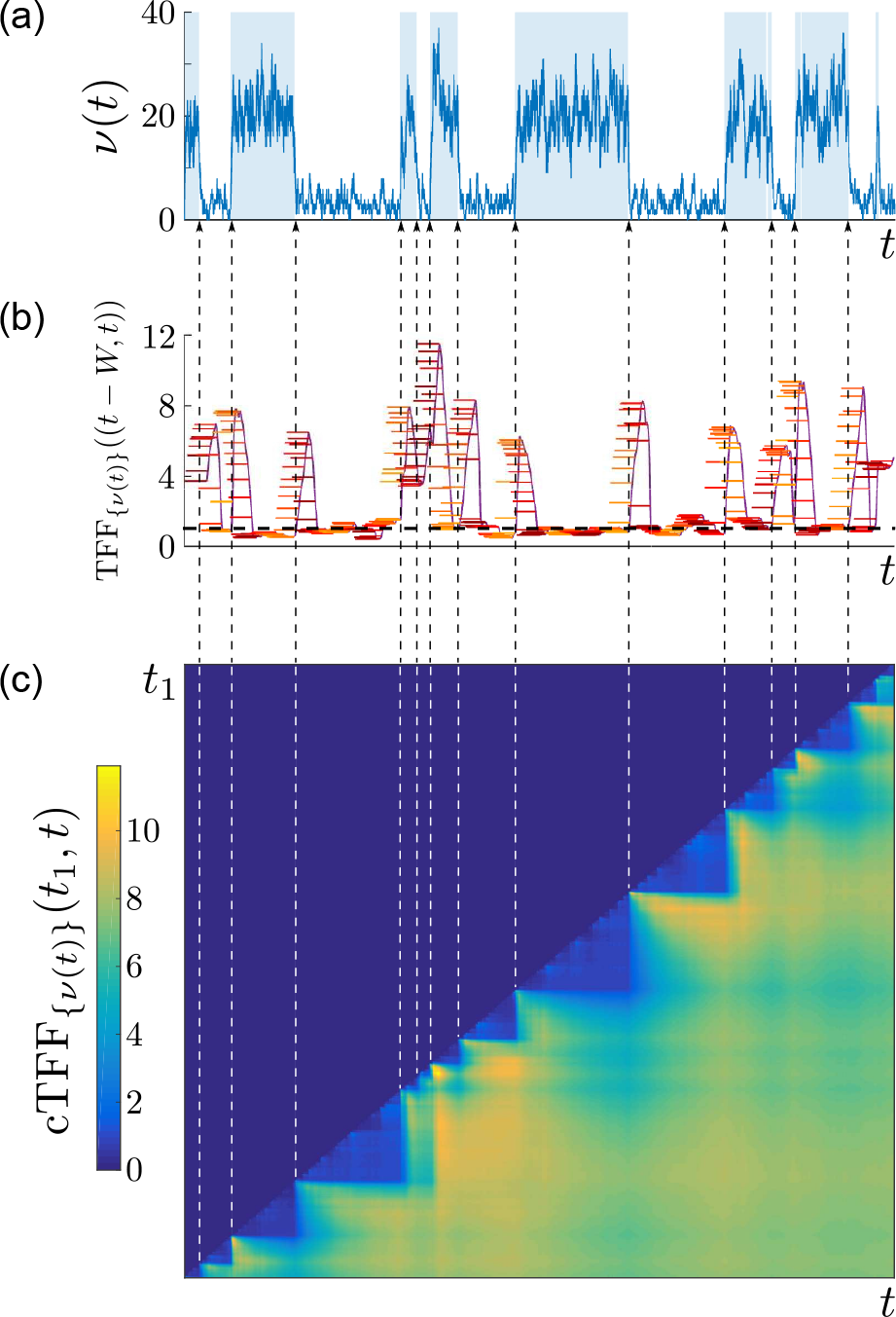
The temporal and cumulative Fano factor. (a) A sample path {*v*(*t*)}_*t*≥0_ of mRNA counts from the (leaky) RT model. The time periods when the gene is in the active state are shaded. (b) The temporal Fano factor (30), TFF_{*ν*(*t*)}_((*t* − *W,T*)), computed over a time window *W* of fixed length indicated by the horizontal bars at each *t*. When *W* extends over a stationary section of the sample path, TFF is close to unity, corresponding to the Poisson distribution (black dashed line). (c) Heatmap of the cumulative Fano factor (31), cTFF_{*v*(*t*)}_(*t*_1_,*t*), defined only for *t* ≥ *t*_1_. Note the marked step pattern corresponding to the switching times, indicated by dashed lines as a guide to the eye.

Our procedure solves the general transcription-degradation model first, and then arrives at its Poisson mixture form (14). Since only the upstream process is model-specific, different models are solved by obtaining the mixing density *f_X_t__* of the upstream process. It is interesting to note that our solution conforms with the “Poisson representation”, which was proposed by Gardiner and Chaturvedi [65, 66] as an ansatz to solutions of the ME. In their representation, *f_X_t__* had support on the complex plane as a means to obtain asymptotic expansions for certain stationary systems [66], and assumptions regarding vanishing boundary conditions were required. In our case, the Poisson mixture (14) is obtained directly as a result of transcription-degradation models, and the mixing density *f_X_t__*, with support on the positive real line, has a clear physical interpretation in terms of single-cell sample paths for a range of time-varying systems.

**FIG. 9.**
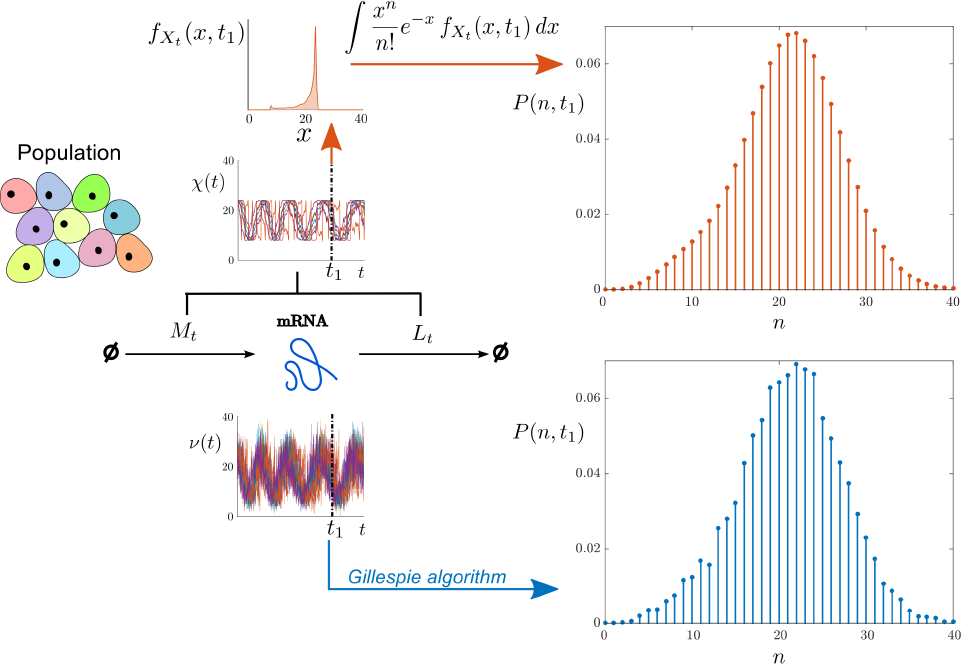
Efficient sampling of the full distribution *P*(*n,t*) for transcription with upstream cellular drives. We consider upstream drives governed by the Kuramoto promoter model (18) for *C* = 10,000 coupled oscillatory cells. Sample paths of *N* are simulated directly with the Gillespie algorithm to approximate *P*(*n, t*) at time *t*_1_ (bottom, blue). Alternatively, sample paths of *X* are used to estimate *f_X_t__*, which is then mixed by performing the numerical integration (14) to obtain *P*(*n,t*) (top, red). The latter sampling through *X* is more regular and far less costly: CPU time via *N* is ≈ 36000 s whereas CPU time via *X* is ≈ 0.1 s.

Our solution confers to us two broad advantages. The first is pragmatic: Since *X_t_* is a continuous random variable satisfying a linear random differential equation, we can draw upon the rich theory and analytical results for *f_X_t__*, even for non-stationary models, or we can use ODE and PDE solvers as further options to solve the differential equation for *f_X_t__*. If simulations are still necessary, sampling *P*(*n,t*|*M, L*) directly using stochastic simulation algorithms becomes computationally expensive, particularly if the upstream processes *M* and *L* are time-varying [67]. Instead, we can sample *f_X_t__* (*x,t*) directly and then obtain the full distribution via numerical integration using (14). This approach leads to a significant reduction in computational cost, as shown in Fig. 9.

Our approach can be used to analyze noise characteristics in conjunction with biological hypotheses. For instance, if measurements of additional cellular variables (e.g., cell cycle) are available, they can be incorporated as a source of variability for gene regulation, with the possibility to test biological hypotheses computationally against experimental data. Conversely, it is possible to discount the Poissonian component from data, so as to fit different promoter models to experimental data and perform model comparison [54].

The second advantageous feature of our approach is conceptual. Studying the natural decoupling of the solution into a discrete, Poisson component and a continuous, mixing component, allows us to derive expressions and properties for both ensemble and temporal moments, extending the concept of ‘intrinsic’/‘extrinsic’ noise to dynamic upstream cellular drives. Importantly, all upstream variability gets effectively imbricated through the upstream effective drive *X*, which can be interpreted in terms of biochemical differential rate equations. This analysis clarifies how upstream fluctuations are combined to affect the probability distribution of the mRNA copy number, providing further intuition of the sources of noise and their characteristics. Indeed, stripping the model down to its extrinsic component *f_X_t__* can provide us with additional understanding of its structure and time scales [54]. Further extensions of our solution could lead to physical interpretations of Gardiner’s complex-valued “Poisson representation” [65, 66], and deeper understanding of models with and without feedback [68]. These extensions will be the subject of future work.

## Footnotes

1 We do not use the term stochastic differential equation (SDE), because SDEs are usually associated with random white noise.

## VII. ACKNOWLEDGMENTS

We are grateful for insightful comments and extended discussions with Juan Kuntz, Philipp Thomas, Martin Hemberg, and Andrew Parry. JD was funded by an EPSRC PhD studentship. MB acknowledges funding from the EPSRC through grants EP/I017267/1 and EP/N014529/1.

## Appendix A: The ‘burn-in’ transient towards stationarity

In Section II A, it was stated that the contribution from the initial condition 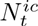 decreases exponentially for biophysically realistic degradation rates {λ(*t*)}_*t*≥0_. As a result, the transcripts that were present at *t* = 0 are expected to degrade in finite time, and the long-term population is expected to be composed only of mRNA molecules that were transcribed since *t* = 0.

Let the initial condition be described by a random variable N0 with a given probability distribution. It follows from Eqs (8)-(9) that

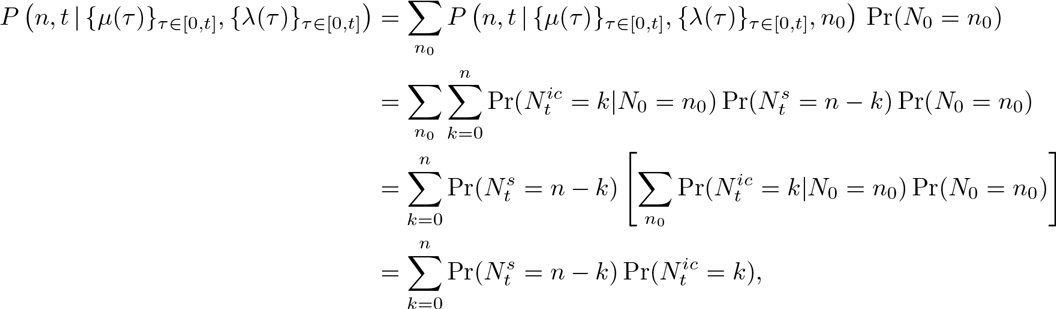

where 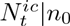 and 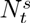 are distributed according to Eqs (5)-(7), and 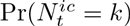 is the mixture of the time-dependent binomial distribution (5) and the distribution of the initial condition *N_0_*.

## 1. ‘Burn-in’ transience in the model with constant transcription and degradation

## The decay towards stationarity

To understand the ‘burn-in’ period more explicitly, consider the simplest example of the gene transcription model (1) with constant transcription and degradation rates *μ* and λ, and assume that there are initially *n*_0_ mRNA transcripts. Given that *N*_0_ ~ *δ*_0,*n*_0__, the solution is given by:

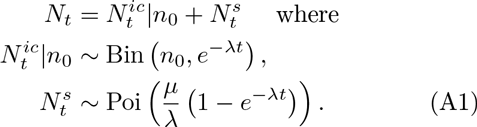

Hence as *t* → ∞, the distribution will tend towards Poi(*μ*/λ), the stationary distribution of the population. This is a well-known result in the literature [66, 69].

## *Starting at stationarity: the time-dependent* 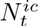 and 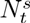 *balance each other at all times*

It is illustrative to consider the dynamics of this system when the initial condition is chosen to be the stationary distribution. In this case, the breakdown of *N_t_* into the time-dependent components 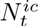 and 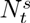 will need to reproduce the stationary distribution at all times *t* > 0, with no ‘burn-in’ period.

To see this,let the initial distribution start at stationarity, i.e., *N*_0_ ~ Poi(*μ*/λ) and

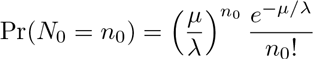

The distribution of 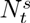 is still given by (A1) and the contribution of 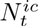 is given by

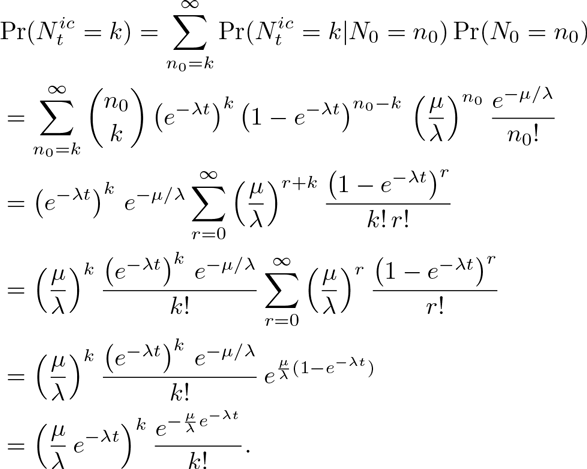

In other words, 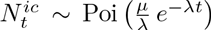, which cancels the (A1) contribution from 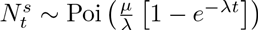. Therefore

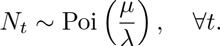

This example shows that 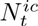 and 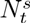 will combine to reproduce a stationary distribution at all times *t* > 0, when the system starts at stationarity so that there is no ‘burn-in’ transient.

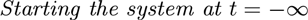

The same is true if the system is not stationary but we start the system at *t* = −∞ with any initial condition. Then, for *t* > 0, the system will be independent of the initial condition and will be described by 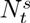.

Let us denote the state of the system for *t* > 0 by the *attracting* distribution *P*_*_. Although 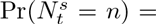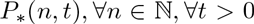, we wish to distinguish *P*_*_ from *P_s_* because we only have equality of the two distributions when the system starts at *t* = −∞. *P*_*_ can be thought ofas an inherent property of the system, analogous to the stable point of a dynamical system that moves in time (sometimes called a *chronotaxic system* [70]).

If Pr(*N*_0_ = *n*_0_) = *P*_*_(*n*_0_, 0) for all *n*_0_ in Eq. (9), the contributions from 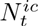 and 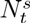 balance each other as they did in the case of stationarity with *N*_0_ ~ Poi(*μ*/λ), and we would have *P*(*n, t*) = *P*_*_(*n, t*), ∀*n* ∈ ℕ, ∀*t* > 0 (recall that the breakdown 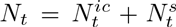 simply resolves the existing mRNA molecules into those that were present at *t* = 0, and those that were transcribed since *t* = 0). Thus we only observe an initial transient period if the initial distribution starts away from its attracting distribution at *t* = 0. In all other cases, the following mathematical formulations are equivalent: *i*) assume that the system was initialised at *t* = −∞ and consider only 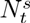, or *ii)* use the initial distribution *P*_*_(*n*, 0) for all *n* at *t* = 0, and consider 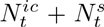.

In this work, we focus on the time dependence of *P*(*n, t*) induced through non-stationarity of the parameters, and/or synchronous behaviour of the cells within the population. Hence, unless otherwise stated, in this work we assume that the system was initialised at *t* = −∞ and that the distribution of 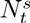 is the attracting distribution *P*_*_(*n,t*) for all *t* > 0, i.e., we neglect the contribution from 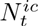.

## 2. ‘Burn-in’ transience in the random telegraph model

A time-dependent solution of the probability generating function for the random telegraph model appeared in Ref. [13], although the explicit expression for *P*(*n, t*) was omitted. As discussed above, the RT model represents asynchronous and stationary behaviour, hence the time-dependence appears only through convergence to the stationary distribution from the initial condition. We include the derivation here for completeness and to complement Fig. 6.

Consider the RT model depicted in Fig. 3a. Assuming that every cell is initialised in the inactive state with *n_0_* = 0 mRNA molecules, the probability generating function for the cell population is [13]:

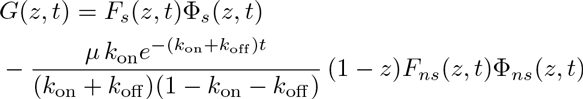

where *F_s_*(*z,t*): = _1_*F*_1_[−*k*_on_, 1 − *k*_on_ − *k*_off_; *μe*^−*t*^(1 − *z*)], Φ_*s*_(*z,t*): = _1_*F*_1_[*k*_on_,*k*_on_ + *k*_off_; *μ*(1 − *z*)], *F_ns_*(*z,t*): = _1_*F*_1_[*k*_off_, 1 + *k*_on_ + *k*_off_; *μe^−t^*(1 − *z*)], and Φ_*ns*_(*z,t*): = _1_*F*_1_[1 − *k*_off_, 2 − *k*_on_ − *k*_off_;−*μ*(1 − *z*)]. Here _1_*F*_1_(*a,b*;*z*) is the confluent hypergeometric function [56].

Using the general Leibniz rule for differentiation, and omitting other details of the differentiation, we obtain

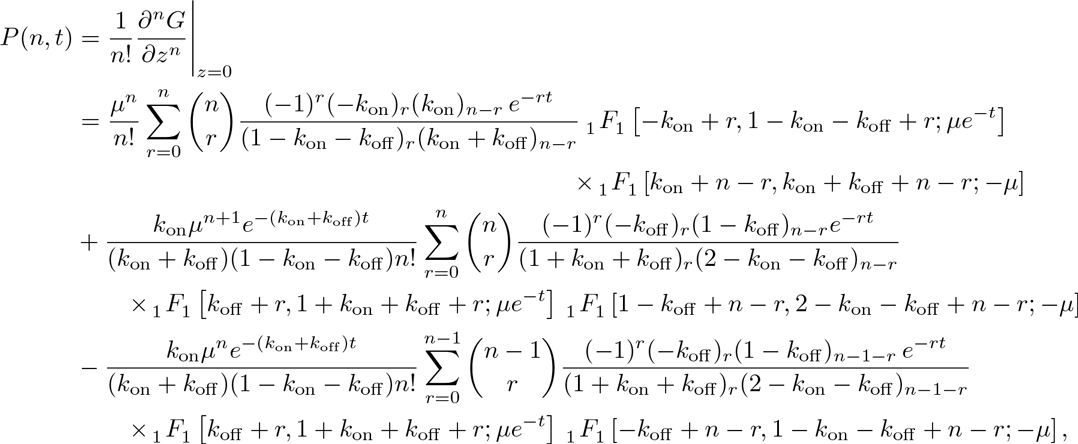

where (*a*)_*m*_: = *a*(*a* + 1)… (*a* + *m* − 1) is Pochhammer’s function [56].

As *ς* → 0, _1_*F*_1_[*a*,*b*;*ς*]. Hence as *t* → ∞,*P*(*n,t*) → *P*(*n*), and we recover the known stationary solution:

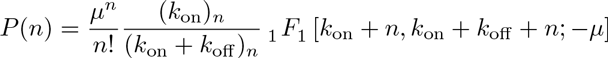

## Appendix B: The stationary solution for the refractory promoter model (3 cyclic promoter states)

As explained in Section IIIB 2, the stationary solution of the 3-state cyclic model describing a refractory promoter is obtained by solving the set of equations:

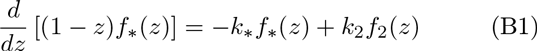

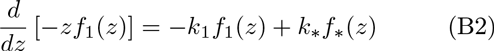

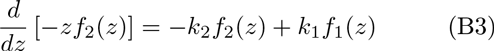

to obtain an expression for *f_Z_* = *f*_*_ + *f*_1_ + *f*_2_.

Note that the transition matrix [*k_sr_*] containing the kinetic constants on the right-hand side of Eqs(B1)-(B3) is singular and hence λ = 0 is an eigenvalue. Furthermore, by Gershgorin’s circle theorem the non-zero eigenvalues of [*k_sr_*] have negative real parts, so a stationary solution always exists, i.e.,the probabilities *π_i_* = *∫*_*x*_*f_i_*(*x*)*dx* of being in state *S_i_* evolve to an equilibrium state given by the eigenvector 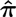 associated with the eigenvalue λ = 0. Note that 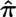 must be normalised so that the elements sum to 1. It can easily be shown that

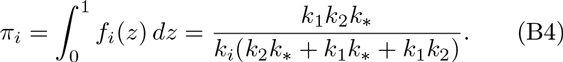

Now, integrating Eqs (B1)-(B3) and using Eq. (B4) we obtain the boundary values *f*_1_(1) = *f*_2_(1) = 0 and *f*_*_(0) = 0. Also, summing Eqs (B1)-(B3) and integrating gives

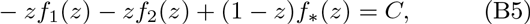

where *C* is a constant. Eq. (B5) is true for all *z* ∈ [0,1], so we can substitute in *z* = 0 or *z* = 1 and use the fact that *f*_1_(1) = 0 and *f*_1_(1) = 0, or *f*_*_(0) = 0, to show that *C* = 0. Hence

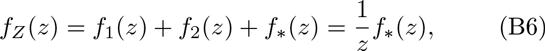

so we need only solve Eqs (B1)-(B3) for *f*_*_(*z*), the marginal probability density corresponding to the active state. Using (B6) and substituting into (B1)-(B3), we then obtain the following equation for *f*_*_(*z*):

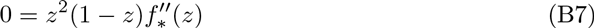

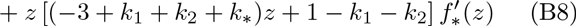

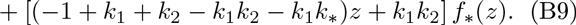

Set *f*(*z*) = *z^c^u*(*z*), where *c* is a constant that we can choose, to transform (B7) into

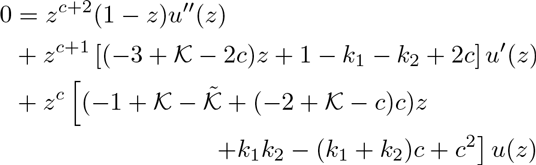

where 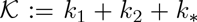 and 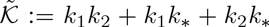. From here, we set the last term on the right hand side to zero by choosing *c* = *k*_1_ or *c* = *k*_2_. We can then divide through by *z*^*c*+1^ to obtain an equation in the form of the hypergeometric equation [56]. For example, for *c* = *k*_1_ we obtain

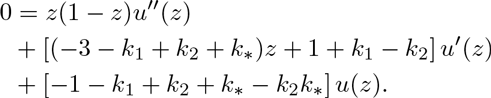

When none of 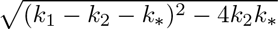, *k*_1_ − *k*_2_ or *k*_*_ are integers, we can write down the solution [56]:

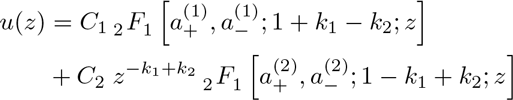

where *C*_1_ and *C*_2_ are constants of integration, _2_*F*_1_ [*a, b*; *c*; *z*] is the Gauss hypergeometric function [56s], and 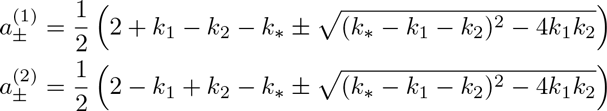.

The solutions for the other cases are similar and are also given in [56]. Hence *f*_*_(*z*) = *z*^*k*_1_^ *u*(*z*) is given by

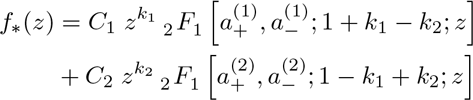

The same expression for *f*_*_(*z*) is obtained if we choose *c* = *k*_2_ instead, so finally we can write down the general solution for *f_Z_*(*z*) = *f*_*_(*z*)/*z*:

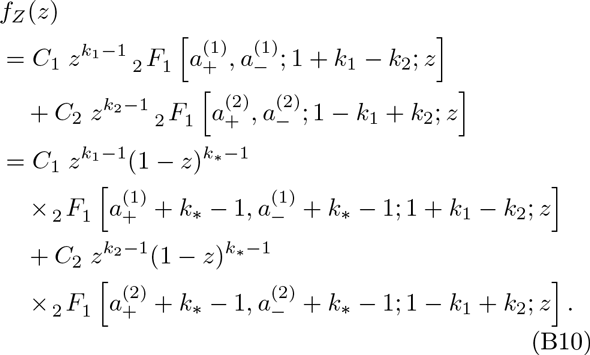

Here, *C*_1_ and *C*_2_ are normalising constants that ensure the integral constraints

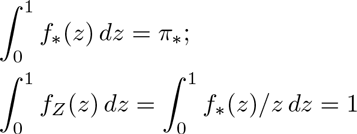

are satisfied. These constants are conveniently obtained from a Mellin transform identity (see Ref. [71], p. 152):

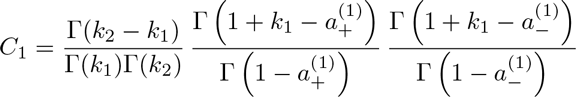

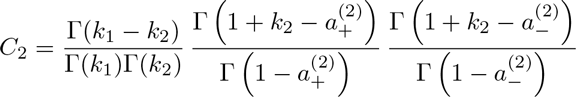

where Γ(.) is the Gamma function [56]. Eq. (B10) is useful for comparisons with the 2-state random telegraph model.

